# Crosstalk between microglia and patient-derived glioblastoma cells inhibit invasion in a three-dimensional gelatin hydrogel model

**DOI:** 10.1101/2020.08.06.238345

**Authors:** Jee-Wei Emily Chen, Jan Lumibao, Sarah Leary, Jann N. Sarkaria, Andrew J. Steelman, H. Rex Gaskins, Brendan A. C. Harley

## Abstract

**Background:** Glioblastoma is the most common and deadly form of primary brain cancer, accounting for more than thirteen thousand new diagnoses annually in the United States alone. Microglia are the innate immune cells within the central nervous system, acting as a front-line defense against injuries and inflammation via a process that involves transformation from a quiescent to an activated phenotype. Crosstalk between GBM cells and microglia represents an important axis to consider in the development of tissue engineering platforms to examine pathophysiological processes underlying GBM progression and therapy.

**Methods:** This work used a brain-mimetic hydrogel system to study patient-derived glioblastoma specimens and their interactions with microglia. Here, glioblastoma cells were either cultured alone in 3D hydrogels or in co-culture with microglia in a manner that allowed secretome-based signaling but prevented direct GBM-microglia contact. Patterns of GBM cell invasion were quantified using a three-dimensional spheroid assay. Secretome and transcriptome (via RNAseq) were used to profile the consequences of GBM-microglia interactions.

**Results:** Microglia displayed an activated phenotype as a result of GBM crosstalk. Three-dimensional migration patterns of patient derived glioblastoma cells showed invasion was significantly decreased in response to microglia paracrine signaling. Potential molecular mechanisms underlying with this phenotype were identified from bioinformatic analysis of secretome and RNAseq data.

**Conclusion:** The data demonstrate a tissue engineered hydrogel platform can be used to investigate crosstalk between immune cells of the tumor microenvironment related to GBM progression. Such multidimensional models may provide valuable insight to inform therapeutic innovations to improve GBM treatment.

## 1. Introduction

Glioblastoma (GBM) is the most common and deadly form of central nervous system cancer [1, 2]. In the United States, it is estimated that more than 13,000 patients are diagnosed with GBM annually. Unlike many other cancers, GBM rarely metastasizes to a secondary organ, but instead diffusely infiltrates throughout the brain. The current standard of care for treating GBM consists of maximal surgical resection, radiotherapy, and concomitant and adjuvant chemotherapy with temozolomide [1, 3, 4]. Despite this aggressive treatment strategy, GBM tumors commonly recur with a median survival of less than 18 months, and fewer than 5% of patients surviving to five years [5-11]. A central focus for improving GBM therapy is developing new tools to understand pathophysiological processes driving GBM invasion of the brain. Improved therapy will likely require both an improved understanding of which cells within the heterogeneous cell population of GBM tumors invade surrounding tissues, and the extent to which cell-cell crosstalk within heterogeneous cell cohorts contribute to GBM invasion and mortality [12-18].

Microglia (MG) are resident immune cells of the CNS [19, 20]. In healthy individuals, microglia constantly survey their surroundings and maintain tissue homeostasis by removing apoptotic cells and promoting neuro-network generation [20-22]. Studies of the GBM tumor microenvironment have demonstrated that infiltrative and resident immune cells, such as microglia, may comprise up to a third of the solid tumor mass [20-22]. Morphologically, quiescent microglia typically exhibit a ramified (branching and elongated) shape. Upon stimulation in response to inflammation, disease, or tumor growth, microglia cell processes become hypertrophic and, in some cases, retract causing the cell to take on an ameboid appearance. While the number and phenotype of immune cells have been associated with patient prognosis [19, 23-27], detailed analysis of crosstalk between GBM cells and microglia are difficult to evaluate in vivo. Thus, there is a need for an experimental platform to rigorously investigate interactions between GBM cells and microglia, as well as to identify factors associated with microglia-GBM crosstalk that may alter GBM cell invasion and therapeutic response. Cancer tissue engineering platforms that integrate biomaterial mimics of the tumor microenvironment with primary cells and biomolecules are increasingly used to investigate pathophysiological processes difficult to examine in vivo [28, 29].

We previously developed a gelatin-based hydrogel model platform to investigate pathophysiological processes underlying GBM cell invasion and therapeutic response. Notably, we observed that biophysical (hyaluronan content and molecular weight) and metabolic (hypoxia) transitions in the GBM tumor microenvironment both significantly alter GBM invasion [30-32]. More recently, we adapted this system to profile cytokine-based crosstalk between cells within the GBM tumor microenvironment, identifying secreted factors generated by an artificial perivascular niche that can accelerate GBM cell invasion [33]. The objective of the present study was to adapt this established hydrogel platform and cytokine analysis protocols to examine the effects of microglia within the GBM tumor microenvironment on GBM gene expression and invasiveness using patient-derived GBM specimens that maintain patient-specific morphologic and molecular phenotypes [34, 35].

## 2. Materials and Methods

### 2.1. Cell culture

#### Human GBM cells

Patient derived GBM cells (PDCs) were obtained from Mayo Clinic (Rochester, Minnesota). All specimens used in this study (GBM12 and GBM39) were derived from tumors from different patients then maintained as patient-derived xenografts in nude mice [34, 35]. All patients consented to the use of their tumor tissue in support of this research, and the use of the patient tissues received prior institutional review board authorization. GBM12 exhibits overexpressed epidermal growth factor receptor (*EGFR*^*OE*^) and displays medium invasive potential in vivo. GBM39 possesses an *EGFR* variant III mutation (*EGFR*^*vIII*^) and displays low invasive potential in vivo [34, 35]. GBM PDCs were established in Dulbecco’s modified eagle medium (DMEM; Gibco, MD) supplemented with 10% fetal bovine serum (FBS; Atlanta Biologicals, Atlanta, GA) and 1% penicillin/streptomycin (Lonza, Basel, Switzerland) at 37°C in a 5% CO_2_ environment. PDCs were shipped by overnight courier and seeded into hydrogel cultures immediately upon arrival.

#### Human microglia cell line

HMC3 microglia cells (ATCC®CRL-3304, ATCC) were cultured in DMEM supplemented with 10% FBS and 1% P/S. Cells were incubated at 37°C in 5% CO_2_ and passaged upon reaching confluence.

#### Primary mouse microglial cultures

All animal care protocols were in accordance with NIH Guidelines for Care and Use of Laboratory Animals and were approved by the University of Illinois Laboratory Animal Care and Use Committee. Both male and female C57BL/6J mice (The Jackson Laboratory, Bar Harbor, ME) were used to obtain primary microglial cultures as described previously [36]. In brief, neonatal (P1–2) mouse pups were decapitated with scissors, the brains were extracted, and the meninges removed under a dissection microscope (Leica, Wetzlar, Germany). Brain tissues were pooled from entire litters, dissociated in Accutase (Thermo Fisher Scientific, Waltham, MA) followed by washing and removal of excess debris. Cells were seeded onto poly-d-lysine-coated (Sigma-Aldrich, St. Louis, MO) T75-flasks. After approximately eight days of culture, the flasks were shaken at 37°C for 1h at 170 rpm in an orbital shaker (Max Q 4000; Thermo Fisher Scientific). The supernatant containing microglia was then collected [36].

### 2.2. Methacrylated Gelatin Hydrogel fabrication and characterization

#### Synthesis and fabrication

Methacrylamide-functinalized gelatin (GelMA) macromers and GelMA hydrogels were synthesized and prepared as previously described [30, 37]. GelMA degree of functionalization was determined by ^1^H NMR spectroscopy (∼50% degree of functionalization).

#### Hydrogel characterization

The compressive modulus of GelMA hydrogels was measured using an Instron 5943 mechanical tester [31]. Hydrogels were tested under unconfined compression at the rate of 0.1 mm/min, with their Young’s modulus obtained from the linear region of the stress-strain curve (2.5%–17.5% strain).

#### Cell number determination

Cell-containing hydrogels were made similarly but with addition of cell suspensions (5000 cells per 25 µL hydrogel) or cell spheroids (5000 cells per spheroid per 25 µL hydrogel) to the pre-polymer solution prior to being placed into Teflon molds (0.2 mm thick, 5 mm radius) and then photopolymerized.

#### Hydrogel identification

To distinguish hydrogels containing PDCs versus microglia, all hydrogel specimens containing microglia were cut into half-disks before being placed into culture while GBM seeded hydrogels were maintained as full disks.

### 2.3. Protein Isolation and Western Blotting

Proteins from HMC3 microglia cultured in hydrogels were isolated using protocols described previously [30-32]. Western blots (2 µg per lane) were probed with primary antibodies specific for CD68 (ab213363, 1/500 in blocking buffer; Abcam, Cambridge, UK) or β-actin (4967S, 1/1000 in blocking buffer; Cell Signaling Technology, Danvers, MA), stained via secondary antibody (7074S, 1/2500 in TBST; Cell Signaling Technology), then imaged via an Image Quant LAS 4010 chemiluminescence imager (GE Healthcare, Chicago, IL). Band intensities were quantified using ImageJ and normalized to β-actin intensities.

### 2.4. Immunofluorescence staining and imaging

Image-iT™ Fixative Solution (Invitrogen, Carlsbad, CA) was used to fix HMC3 microglia seeded hydrogels. Cells were permeabilized with 0.1% Triton X-100 (Sigma-Aldrich) in PBS and stained with Alexa Fluor™ 488 Phalloidin (Invitrogen) for F-actin and Hoechst 33342 (Invitrogen) for nuclei following the manufacturer’s protocol. Stained samples were imaged using a Zeiss LSM 700 confocal microscope.

### 2.5. RNA extraction and quality analysis

Total RNA was extracted using an RNeasy Plant Mini Kit (Qiagen, Hilden, Germany) with an additional step using an RNase-free DNase set (Qiagen) for DNase digestion. RNA integrity number (RIN) was determined using an Agilent 2100 bioanalyzer with all samples exhibiting a RIN > 8.

### 2.6. RNAseq analysis

Libraries for RNA sequencing were prepared using the TruSeq Stranded mRNAseq Sample Prep Kit (Illumina, San Diego, CA). The libraries were quantitated by qPCR and sequenced on one lane for 101 cycles from one end of the fragments on a NovaSeq 6000 (Illumina) using a NovaSeq SP reagent kit and yielded 400 to 500 million single reads per lane. Library quality check was done using FastQC (version 0.11.8). Salmon version 0.13.1 was used to quasi-map reads to the transcriptome and quantify the abundance of each transcript. Filtering was set with the threshold of 0.5 counts per million and resulted in 15,901 genes to be analyzed for differential expression that contained > 99.5% of the reads. After filtering, trimmed mean of M values (TMM) normalization in the edgeR package was performed and log2-based count per million values (logCPM) were calculated [38-40]. Differential gene expression analysis was performed using the limma-trend method. Multiple testing correction was done using the False Discovery Rate (FDR) method [41-43]. Functional annotation was performed, gene ontology (GO) KEGG pathways were identified using an overrepresentation test [44]. The 50 genes with the highest fold-changes were subsequently examined in Cytoscape using the application iRegulon to predict transcriptional regulators [45, 46]. Putative transcription factors or motifs with a normalized enrichment score larger than three (NES > 3) is considered to be potential regulators.

### 2.7. Spheroid invasion assay

GBM12 PDCs were counted and resuspended into 5000 cells/200 µL media per well and distributed to Corning spheroid microplates. Plates were centrifuged for 100*g* for 1 minute to assist spheroid formation then placed into incubator (37°C, 5% CO_2_) for 24 hours. Plates were then incubated for additional 24 hours with horizontal shaking at 60 rpm. Spheroids were then transferred and mixed with pre-polymerized GelMA solution and photopolymerizated into hydrogels. Spheroid images were acquired using a Leica DMI 400B florescence microscope (Leica, Germany) at days 0 (immediately upon seeding), 1, 3, 5, and 7. Invasion was then quantified via ImageJ and invasion distance reported as fold change of their average radius compared to day 0 as described previously [30, 31].

### 2.8. Quantification of cell number

Cell proliferation was determined using a commercial Vybrant® MTT Cell Proliferation Assay Kit (Invitrogen) adapted from the manufacturer’s protocol as described previously [30, 32, 47].

### 2.9. Secretome profiling

Cell culture media was collected from the following group: GBM-MG co-cultures (co-culture), GBM cells alone (GBM single), or MG cells alone (MG single). The media were spun down (300 × g, 10 minutes) to remove any debris. As an additional control, we created a 1:1 mixture of GBM single and MG single media (Mix) that would not account for any GBM-MG crosstalk mediated shifts in secretome. The secretome for each specimen was profiled using a Proteome Profiler™ Human Angiogenesis Array Kit (Ary007, R&D Systems, Minneapolis, MN) following the manufacturer’s protocol. Blots were imaged using an Image Quant LAS 4010 chemiluminescence imager (GE Healthcare). Dot intensities were quantified with the ImageJ macros toolset Protein Array Analyzer (https://imagej.nih.gov/ij/macros/toolsets/Protein%20Array%20Analyzer.txt, Gilles Carpentier) (Table S1). Data was first normalized by dividing the pixel intensity for each blot by the average positive control pixel intensity (on each membrane). We calculated intensity fold change between co-culture and mix groups, identifying factors that displayed a larger than 1.5-fold change; targets showing >0.75-fold change relative to positive reference spot intensities was plotted.

### 2.10. Statistical Analysis

Statistical analysis for Western blot was performed via t-test. Analysis of MTT was performed via one-way ANOVA followed by Tukey’s HSD post-hoc test. Analyses of invasion was performed via two-way ANOVA followed by Tukey’s HSD post-hoc test. Significance level was set at p<0.05 unless stated otherwise (p<0.01 and p<0.0001). A minimum of n = 3 samples was used for all analyses and specified in each result section. Error bars are plotted as the standard error.

## 3. Results

### 3.1. Co-culture system assembly and mechanical testing of GelMA hydrogel

GBM and MG cells were maintained in three-dimensional culture in a 4 wt% methacrylamide-functinalized gelatin (GelMA) that exhibited a physiologically relevant Young’s modulus (1.04±0.10 kPa; n=14; Figure 1A) [48, 49]. Individual (*GBM single, MG single*) cultures were generated as either GBM or MG-seeded hydrogel disks in separate culture wells. Alternatively, GBM and MG seeded hydrogel disks were cultured together in the same well (*co-culture*), allowing exchange of secreted factors between hydrogel disks but not allowing direct cell-cell contact (Figure 1B).

**Figure 1.**
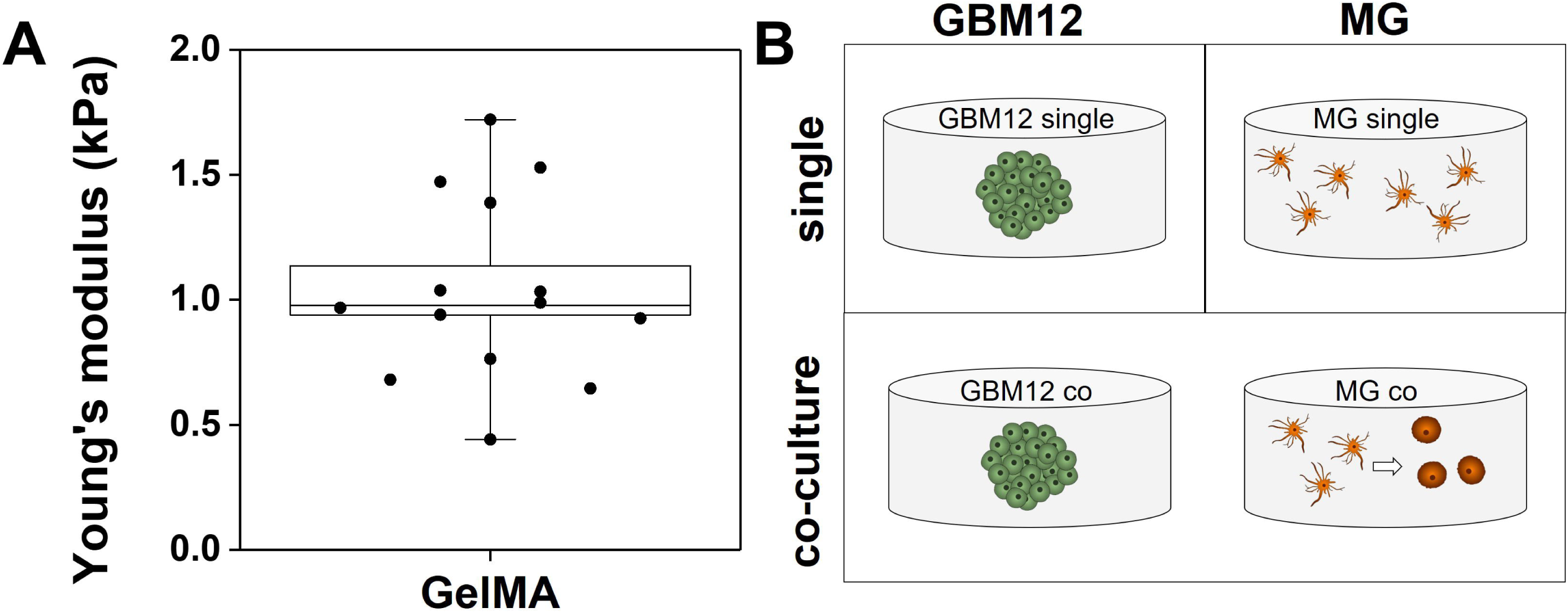
GelMA hydrogel biophysical parameters and experimental schematic. (A) Mechanical testing results of GelMA hydrogel confirmed its physiologically relevant Young’s modulus (1.04±0.10 kPa; n=14). (B) Schematic representation of the culture system, that either has a single type of cell-seeded hydrogel disk per well (*GBM single, MG single*) or contains two distinct hydrogel disks (GBM-MG *co-culture*). GBM-MG *co-culture* allows soluble signaling without physical GBM-MG contact.

### 3.2. Microglia become activated when co-cultured with GBM cells

Microglia displayed significant morphological changes as a result of GBM-MG crosstalk (Figure 2). HMC3 microglia cultured individually in GelMA hydrogels (*MG single*) exhibited the characteristic elongated shape of quiescent MG. However, in response to co-culture with GBM12-seeded hydrogels (GBM-MG *co-culture*), MG adopted a rounded ameboid shape associated with activation. To confirm this shift in activation status, changes in the expression of the lysosomal protein CD68 were examined via Western blot (n=3). While CD68 is normally expressed at low levels in quiescent microglia, CD68 expression was increased in microglia co-cultured with GBM cells.

**Figure 2.**
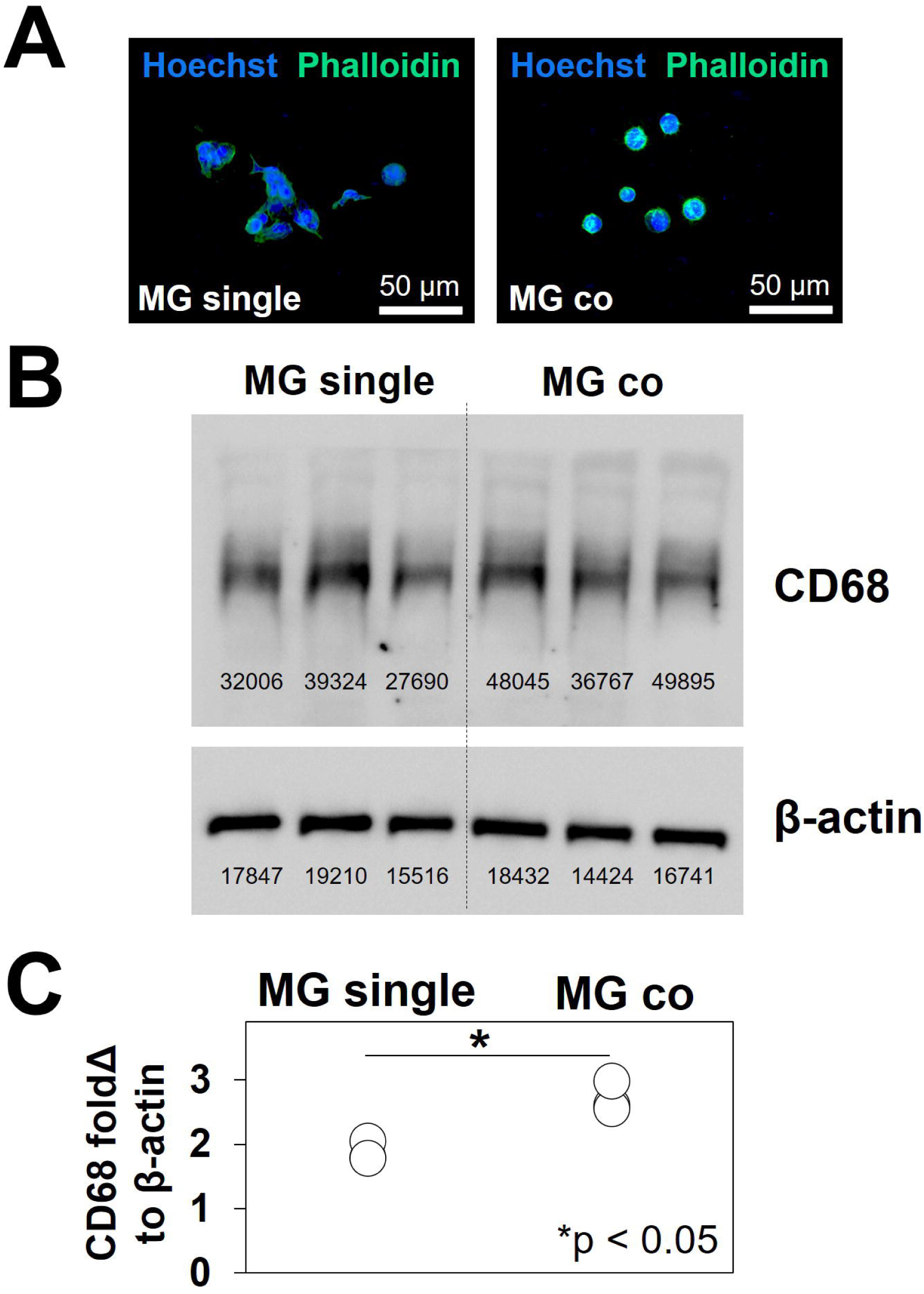
Verification of microglia activation via morphological and protein expression analyses. (A) Immunofluorescence staining of Alexa488Phalloidin^+^ actin (green) showing HMC3 microglia morphology in *MG single* (elongated quiescent) or GBM-MG *co-culture* (ameboid, activated). Also, Hoechst 33342^+^ nuclei stained blue. (B) Western blot staining CD68 and β-actin in triplicate. MG single on the left and MG co-culture on the right. (C) CD68 protein expression levels (n=3) in microglia increased as a result of GBM co-culture. * p<0.05.

### 3.3. Soluble factors produced by microglia altered GBM transcriptomic profiles

RNAseq analysis was used to examine global shifts in the transcriptomic profile of GBM12 specimens as a result of MG crosstalk. Two RNAseq datasets were compared: (1) a *co-culture specimen* containing RNA isolated from the GBM-seeded hydrogels kept in physically separated co-culture with HMC3 MG-seeded hydrogels; and (2) a *GBM single* culture specimen containing RNA isolated from GBM-seeded hydrogels cultured independently (n=3 for each group). This comparison allowed investigation of global shifts in GBM gene expression profiles only due to GBM-MG crosstalk (Figure 3). Differential gene expression (DE) analysis was performed using an FDR-adjusted p-value set to□0.05. In total, there were 3409 DE genes, of which 1563 were up-regulated and 1846 were down-regulated as a result of co-culturing GBM cells with microglia. Gene ontology (GO) analysis for biological process ontology showed upregulated genes were associated with proliferation and RNA/DNA replication and proliferation, while down regulated genes were associated with motility, adhesion, and invasion (Supplementary Table S1; Figure 3B).

**Figure 3.**
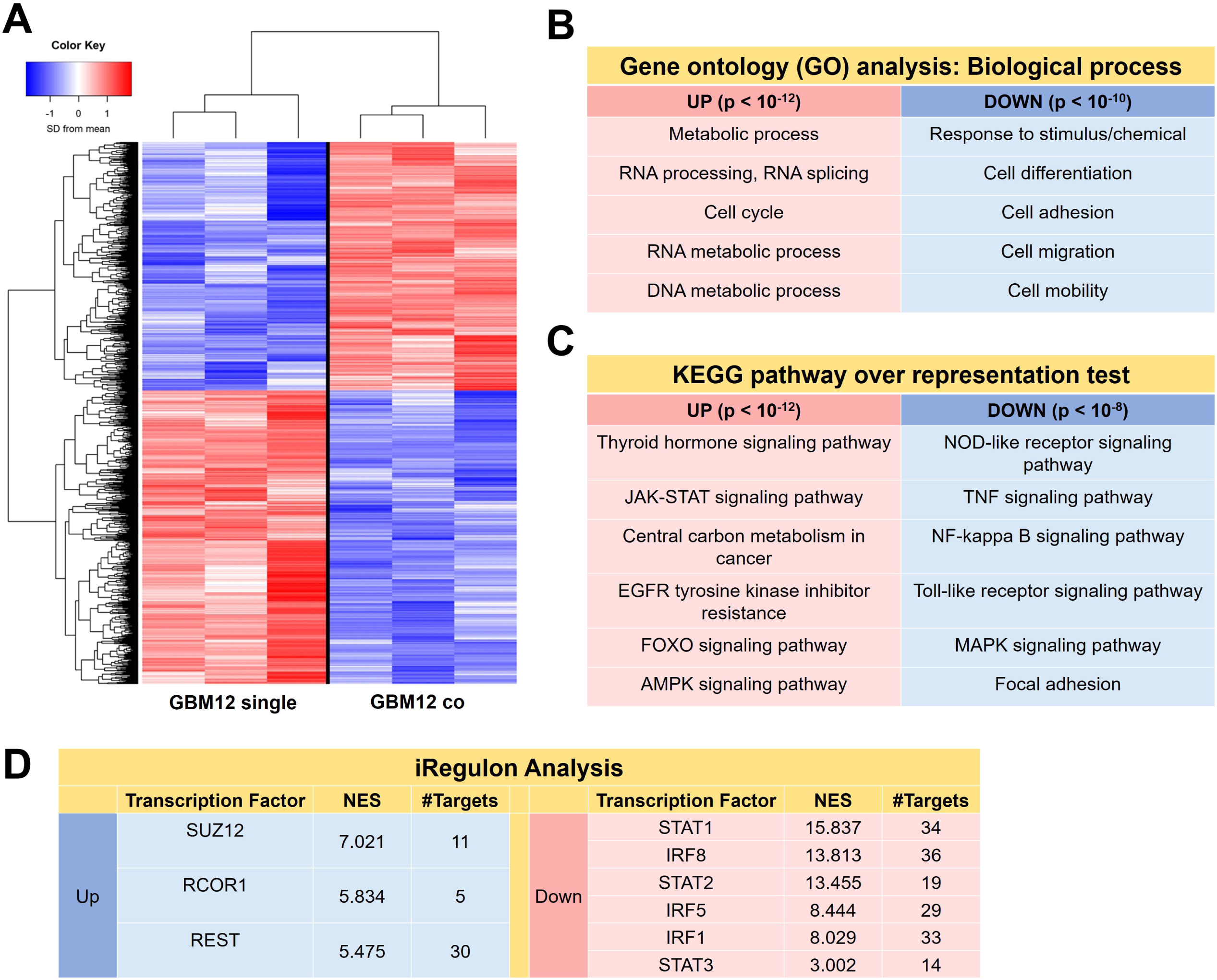
Soluble signals from microglia induced shifts in GBM12 cell transcriptome. (A) RNAseq and differential gene expression analysis (p<0.05) identified 1563 upregulated genes and 1846 downregulated genes in GBM12 co compared to GBM12 single. Selected (B) gene ontology (GO) and (C) KEGG pathway terms suggest GBM-MG interactions underlie upregulated expression of genes involved in GBM proliferation and genes with diminished expression associated with GBM invasion. (n=3 for GBM12 single and GBM12 co) (D) Significant putative genes from the top 50 genes with normalized enrichment score > 3 were identified via iRegulon.

KEGG pathway analysis was used to identify the top 20 up- and down-regulated pathways (Table 2; Figure 3C). GBM-MG *co-culture* downregulated multiple immune response-related pathways in GBM cells including nucleotide-binding and oligomerization domain (*NOD*)-like receptors (*NLRs*), tumor necrosis factor (*TNF*) signaling, nuclear factor kappa-beta (*NF-κB*) signaling, toll-like receptor signaling (*TLRs*), and mitogen-activated protein kinase (*MAPK*) signaling. GBM-MG co-culture decreased focal adhesion pathway activation. Upregulated KEGG pathways included those involved in cell proliferation and survival included thyroid hormone signaling, Janus kinase-signal transducer and activator of transcription pathway (*JAK-STAT*) signaling, central carbon metabolism, forkhead box transcription factors (*FOXO*) signaling, and adenosine monophosphate-activated protein kinase (*AMPK*) signaling. Intriguingly, *EGFR* tyrosine kinase inhibitor resistance was also increased, indicating that the co-culture might also alter the therapeutic response to classes of drugs targeting tyrosine kinase inhibitors. iRegulon analysis of upregulated genes showed high normalized enrichment scores for *SUZ12* (NES=7.021), *RCOR1* (NES=5.834) and *REST* (NES=5.475). iRegulon analysis also showed high normalized enrichment scores for downregulated genes *STAT1/2/3* (NES=15.831/13.455/3.002) and *IRF1/5/8* (NES=8.029, 8.440, 13.813). Together, these analyses indicate that co-culture with microglia induces significant shifts in GBM transcriptome linked to increased proliferative behavior and decreased invasive potential.

### 3.4. Microglial soluble factors increased proliferation but inhibited GBM invasion

GBM proliferation and invasion was subsequently examined as a function of MG co-culture. GBM12 cell proliferation was significantly increased in response to GBM-MG co-culture (Figure 4A). MG *co-culture* strongly inhibited GBM12 invasion over the course of a seven-day spheroid-based invasion assay, with effects observed as early as after 24-hours (Figure 4B,C). A consistent inhibitory effect of MG co-culture on GBM invasion was observed as well for a different source of microglia (primary neonatal microglia, nMG) and a different patient derived GBM cell population (GBM39: *EGFR*^*VIII*^). GBM39 show reduced invasive potential in vivo compared to GBM12, but nonetheless GBM39 invasion was again significantly inhibited in the multi-dimensional hydrogel model in response to nMG co-culture, with effects observed as early as day 3 (Figure 5).

**Figure 4.**
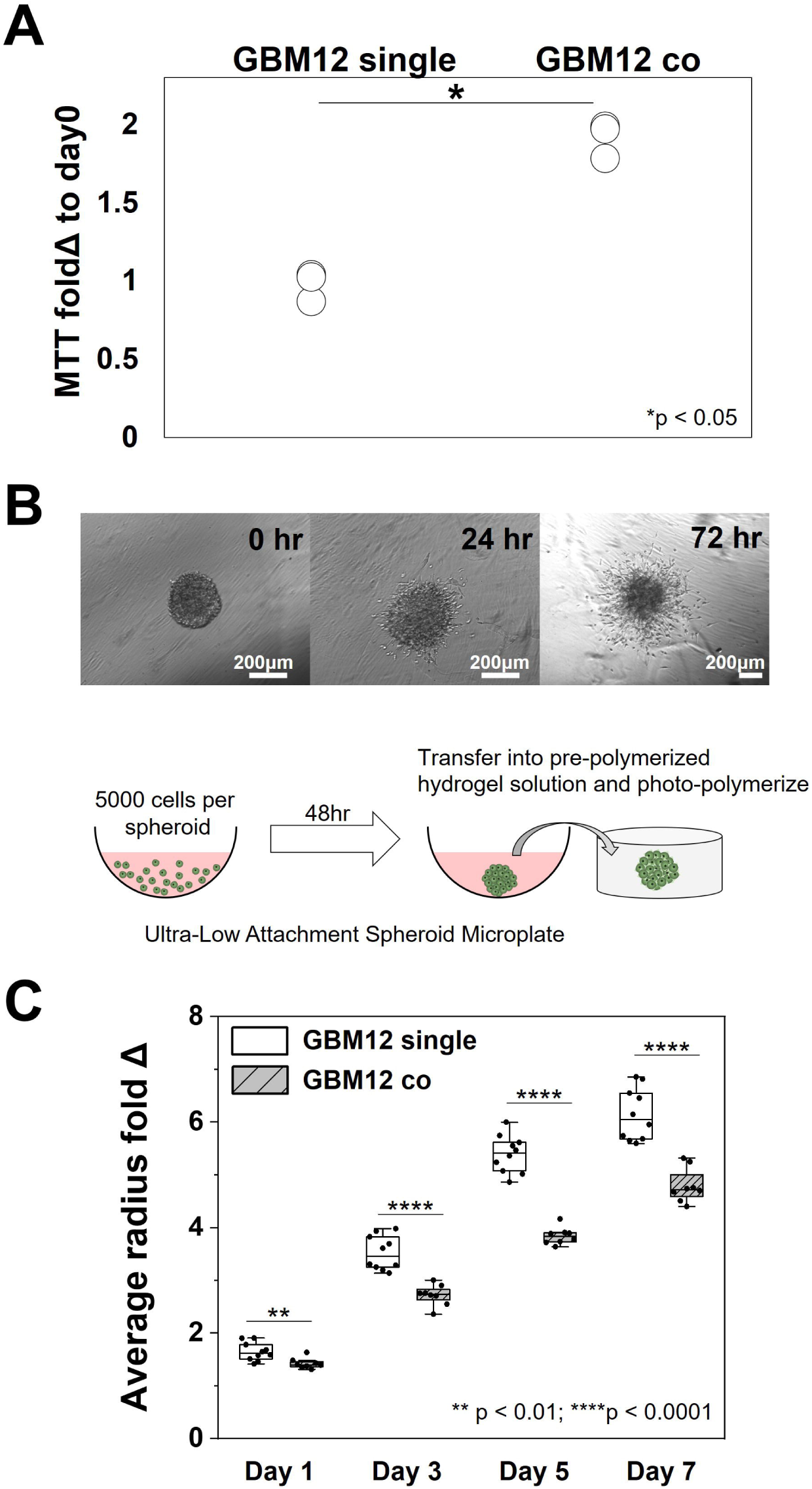
Soluble factors from microglia increase GBM proliferation but significantly inhibit GBM12 invasion. (A) Significant increase in MTT fold change at day 3 in GBM12 cells co-cultured with MG compared to GBM12 single cultures (both n=3). (B) Representative images of spheroid-based invasion assay, with invasion metrics shown as fold-change of average radius of cell spreading compared to day 0 (immediately after culture). (C) GBM12 cells exhibited significantly decreased invasion (n=8) in response to co-culture with HMC3 microglia (vs. GBM12 alone; n=10). *p<0.05. **p<0.01. ****p<0.0001.

**Figure 5.**
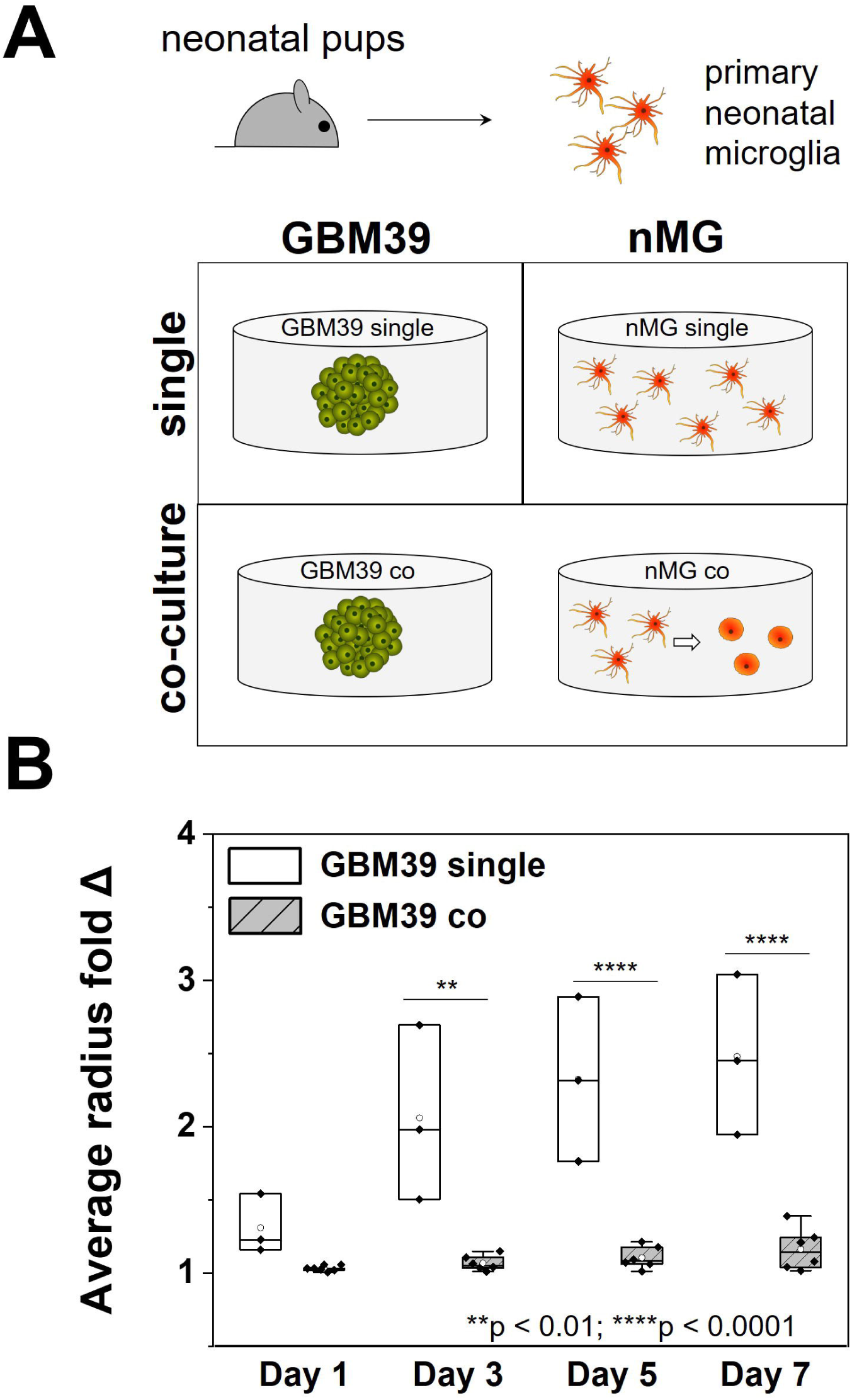
Primary neonatal microglia strongly inhibit GBM invasion. (A) Primary neonatal microglia cells (nMG) were obtained from neonatal mouse pups. nMG and GBM39 seeded hydrogel disks were then either cultured alone (single) or together (co) in the same well of a 24-well plate. (B) GBM39 invasion was significantly decreased due to co-culture with nMG (n=6) compared to GBM39 alone (n=3). **p<0.01. ****p<0.0001.

### 3.5. Profiling MG-GBM secretome using cytokine array

A secretome screen was performed to compare cell culture media from GBM-MG *co-culture* versus GBM or MG monocultures (*GBM single, MG single*) and a *Mix* media (1:1 mixture of *GBM single* and *MG single* medias; Figure 6). Eight factors exhibited a >1.5 fold change in *co-culture* vs. single cultures: chemokine ligand 2 and 3 (CCL2, CCL3); insulin-like growth factor binding protein 3 (IGFBP-3); angiogenin (ANG); heparin-binding epidermal growth factor-like growth factor (HB-EGF); dipeptidyl peptidase-4 (DPP4); Serpin F1; and coagulation factor III (F3). Six factors were expressed at levels greater than 0.75 of the positive reference intensity value within each secretome array (Figure 6C). Of these, CCL2 showed the largest activation (>2-fold change) for GBM-MG *co-culture* versus the *Mix* media control. Granulocyte-macrophage colony-stimulating factor (GM-CSF, or CSF2) and pentraxin-related protein (PTX3) were both highly expressed in cultures containing MG (*MG-single, Mix, co-culture*) media but lowly expressed in *GBM-single* hydrogels. Serpin E1, tissue inhibitor of metalloproteinase-1 (TIMP-1) and vascular endothelial growth factor (VEGF) were expressed across all culture conditions.

**Figure 6.**
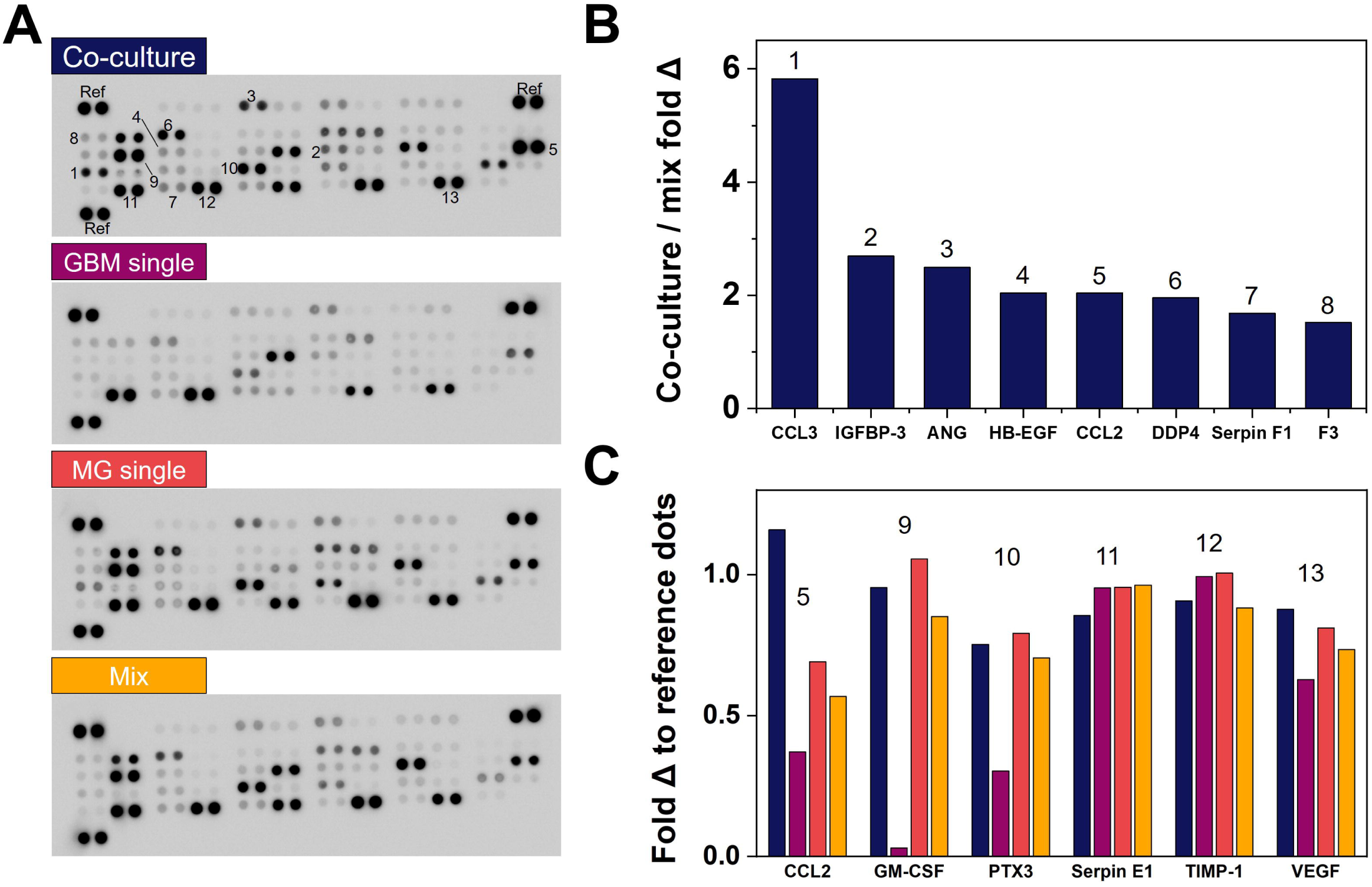
Profiling GBM-MG (GBM12-HMC3) secretome linked to degreased GBM cell invasion. (A) Proteome Profiler™ Array analysis of secretome profiles from conditioned media of 4 distinct culture conditions: GBM-MG *co-culture*; *GBM single*; *MG single*; 1:1 *Mix* of *GBM single* and *MG single*. (B) Eight factors that displayed >1.5 fold change in GBM-MG *co-culture* versus *Mix* media groups. (C) Six factors (from GBM-MG *co-culture*) that displayed greater than 0.75 fold change versus the positive reference dots. Numbers (1-13) correspond to positions labeled on blots in (A).

## 4. Discussion

Cellular crosstalk within the tumor microenvironment provides a powerful avenue of interaction that may significantly shape disease progression. Tools to interrogate cellular crosstalk offer an opportunity to identify novel therapeutic compounds to improve treatment of glioblastoma. This work demonstrates the use of a tissue engineering platform to investigate the role of crosstalk between patient derived GBM specimens and microglia on shifts in the phenotypic, proteomic, and transcriptomic signatures of GBM. GBM-MG crosstalk is bidirectional and induces microglia activation along with shifts in GBM cell activity consistent with the go-or-grow phenomenon [50].

While microglia display a quiescent phenotype in single culture, hallmarks of microglial activation were observed via <u>both</u> morphological changes and increased CD68 expression as a result of GBM-MG co-culture. These results are consistent with hallmarks of microglial activation seen in cases of disease and histopathological analysis of GBM tumors [24, 51]. The nature of this tissue engineering platform allows significant post-culture analysis of GBM cell activity at functional (invasion, proliferation), transcriptomic (RNAseq), and secretomic levels.

While advanced sequencing techniques such as RNA sequencing offer the opportunity to define the transcriptomic signature of cells to aid treatment planning and outcome prediction [39, 42, 43, 52], the design of the hydrogel culture system reported here enabled analysis of shifts in the transcriptome of individual cell populations as a result of heterotypic cell (GBM, MG) crosstalk. GO analyses revealed GBM-MG *co-culture* upregulated genes in a patient-derived GBM specimen associated with cell cycle, RNA/DNA division and metabolic activity. However, genes involved in cell adhesion/migration showed significant downregulation as a result of GBM-MG *co-culture*. These findings indicate tradeoffs in GBM proliferation versus invasion due to MG crosstalk consistent with the go-or-grow dichotomy of GBM cells [32, 50, 53, 54]. Significant decreases were observed in expression of genes associated with *NLR, TNF, NF-κB, MAPK*, and *TLR* pathways in GBM specimens in response to MG *co-culture. NLR* and *TLR* signaling pathways are involved in pathophysiological responses to inflammation and tumor progression [55-57]. Of these, the *NF-κB* signaling pathway is known to be sensitive to *TNF* signaling [58-60] which plays a major role in immune activation [61, 62], breast cancer invasion [63], and driving *TLR* and *MAPK* signaling involved in cell migration and tumor invasion These pathways contribute to heightened immune responsiveness and are involved in angiogenesis and cell migration [55, 56, 58-60], suggesting GBM-MG interactions may inhibit GBM invasiveness.

KEGG analysis also showed strong upregulation in *TH* and *STAT3* signaling, indicating that secreted factors from microglia may promote GBM proliferation, reduce apoptosis, and enhance chemotherapeutic resistance [64-66]. Recently, our group showed STAT3 is strongly activated in GBM, and inhibiting STAT3 can reduce GBM cell proliferation [67, 68]. More, GBM-MG *co-culture* upregulated *FOXO* signaling, which has been linked to therapeutic resistance due to its contribution to DNA repair as well as mediation of oxidative stress. iRegulon analysis showed GBM-MG crosstalk increased enrichment for *SUZ12*, previously shown to be increased in high grade astrocytoma and involved in pathways that regulate glioma proliferation and metastasis [69, 70]. iRegulon analysis also showed GBM-MG crosstalk increased *REST* and *RCOR1*, known to regulate the oncogenic properties of GBM stem cells [71] that associate with therapeutic resistance and recurrence [72, 73]. The *IRF* family has been shown to be significant tumor suppressors, inhibit tumor proliferation and loss of *IRF* genes may contribute to tumor metastasis and invasion [74, 75]. Together, analysis of transcriptomic data support the functional responses of increased proliferation but decreased invasion for GBM cells as a result of GBM-MG interactions [32, 54, 76-78].

The hydrogel platform was subsequently used to experimentally interrogate the influence of GBM-MG crosstalk on GBM proliferative and invasive phenotypes in patient-derived GBM12 cells. GBM12 cells exhibited significantly increased proliferation and significantly inhibited invasion in response to MG co-culture. Strikingly, MG-induced inhibition of GBM12 invasion was observed for multiple combinations of patient-derived GBM specimens and microglia: *EGFR*^*OE*^ GBM12 cells co-cultured with HMC3 microglia and *EGFR*^*vIII*^ GBM39 cells co-cultured with primary mouse neonatal microglia.

Analysis of the combined GBM-MG secretome revealed multiple targets driving the observed shifts in functional and transcriptomic activity. CCL2 and CCL3 are associated with monocyte and macrophage recruitment [79-81] and may act as chemoattractant [79]. Of these, further study of the role of CCL2 in GBM invasion may be particularly warranted, as expression levels were not only significantly increased in GBM-MG *co-culture* (vs. *GBM-single* or *MG-single* cultures) but also compared to the *Mix* control, consistent with synergistic activation of CCL2 secretion due to GBM-MG crosstalk. IGFBP-3, known to regulate cell proliferation, was also increased in GBM-MG crosstalk, though its role in cancer progression remains to be fully understood [82-84]. DPP4 (plasma membrane protein that contributes to immune and metabolic regulation [85, 86]) and HB-EGF (cell metabolic activity and tumor suppression in other cancers [85-87]) were also strongly upregulated in GBM-MG *co-culture*, as was ANG, well-known for its role in angiogenesis and cell proliferation [88, 89], and Serpin F1, known as for its role in suppression of tumor growth and prostate cancer metastasis [90-93]. Previous study by Shinozaki et al.[94] also indicated that cytokines produced by microglia could potentially drive astrocytes towards a neuroprotective phenotype upon brain injuries. However, further analysis is needed to more fully investigate the potential mediators of GBM invasion that arise from GBM-MG crosstalk.

The combination of increased proliferation but decreased invasion aligns with the go-or-grow hypothesis [53], but more importantly demonstrates that crosstalk between MG and GBM cells in the tumor microenvironment may have powerful effects on GBM activities tied directly to tumor progression and patient survival.

## 5. Conclusion

This study describes a tissue engineering platform to examine the role of GBM-microglia crosstalk on processes associated the GBM progression. It also shows the ability to use bioinformatic tools to identify transcriptomic shifts underlying these responses. We show dynamic, two-way interactions between patient derived GBM cells and microglia via paracrine signaling influence both microglia and GBM cell phenotype. Microglia in the presence of patient-derived GBM cells showed morphological shifts associated with activation. Microglia co-culture significantly inhibited GBM invasion but enhanced proliferation that could be captures via three-dimensional spheroid invasion assays and transcriptomic analyses. Future efforts will seek to understand the contribution of GBM-microglia crosstalk on tumor resistance to therapeutics, to reveal candidate signaling axes for rational combinatorial targeting.

## Declarations

### Ethics approval and consent to participate

#### Patient-derived xenograft models

Accrual of patient tumor samples to originally develop these models was performed with approval by the Mayo Clinic Institutional Review Board. The xenograft models being used in this project are derived from patients who are now deceased and therefore do not constitute human subjects research as per NIH guidelines.

#### Primary mouse microglia

All animal care protocols were in accordance with NIH Guidelines for Care and Use of Laboratory Animals and were approved by the University of Illinois Laboratory Animal Care and Use Committee and by the Mayo Clinic Institutional Animal Care and Use Committee.

### Consent for publication

All authors of the manuscript have read and agreed to its content and are accountable for all aspects of the accuracy and integrity of the manuscript.

### Availability of data and materials

Data and metadata associated with this manuscript are available on request to the corresponding author.

### Authors’ contributions

JEC, JL, HRG, AJS, and BACH conceived and designed the experiments. JEC, JL, and SL performed all experiments. JNS provided critical research materials. JEC, JL, HRG, AJS, and BACH analyzed and discussed the data. JEC and BACH. wrote the manuscript. JEC, JL, JNS, HRG, AJS, and BACH edited and revised the manuscript.

## Acknowledgements

We acknowledge the assistance of Dr. Alvaro Hernandez, Dr. Jenny Drnevich and Chris Wright from the Roy J. Carver Biotechnology Center at the University of Illinois Urbana-Champaign for their assistance in RNA quality testing, library preparation, RNA sequencing and analysis. We acknowledge Haw-Wen Hsiao for generating the customized MATLAB code for mechanical testing analysis.

## Abbreviations

AMPK: Adenosine monophosphate-activated protein kinase
ANG: Angiogenin
CCL: Chemokine ligand
CNS: Central nervous system
DE: Differential gene expression
DPP4: Dipeptidyl peptidase-4
EGFR: Epidermal growth factor receptor
F3: Coagulation factor III
FDR: False discovery rate
FOXO: Forkhead box transcription factors
GBM: Glioblastoma
GelMA: Gelatin methacrylate
GM-CSF: Granulocyte-macrophage colony-stimulating factor
GO: Gene ontology
HB-EGF: Heparin-binding epidermal growth factor-like growth factor
IGFBP: Insulin-like growth factor binding protein
IRF: Interferon regulatory factor
JAK: Janus kinase
KEGG: Kyoto Encyclopedia of Genes and Genomes
MAPK: Mitogen-activated protein kinase
MG: Microglia
NES: Normalized enrichment score
NLRs: Nucleotide-binding and oligomerization domain-like receptors
NF-κB: Nuclear factor kappa-beta
PDC: Patient derived GBM cell
PTX3: Pentraxin-related protein
RIN: RNA integrity number
RNAseq: RNA sequencing
STAT: Signal transducer and activator of transcription
TIMP: Tissue inhibitor of metalloproteinase
TLRs: Toll-like receptor signaling
TNF: Tumor necrosis factor
VEGF: Vascular endothelial growth factor

