## supplemental file for "Crosstalk between microglia and patient-derived glioblastoma cells inhibit invasion in a three-dimensional gelatin hydrogel model"

### Supplementary Figures

### Figure S1.

Raw images for β-actin Western blot samples (left three: MG single; right three: MG co)

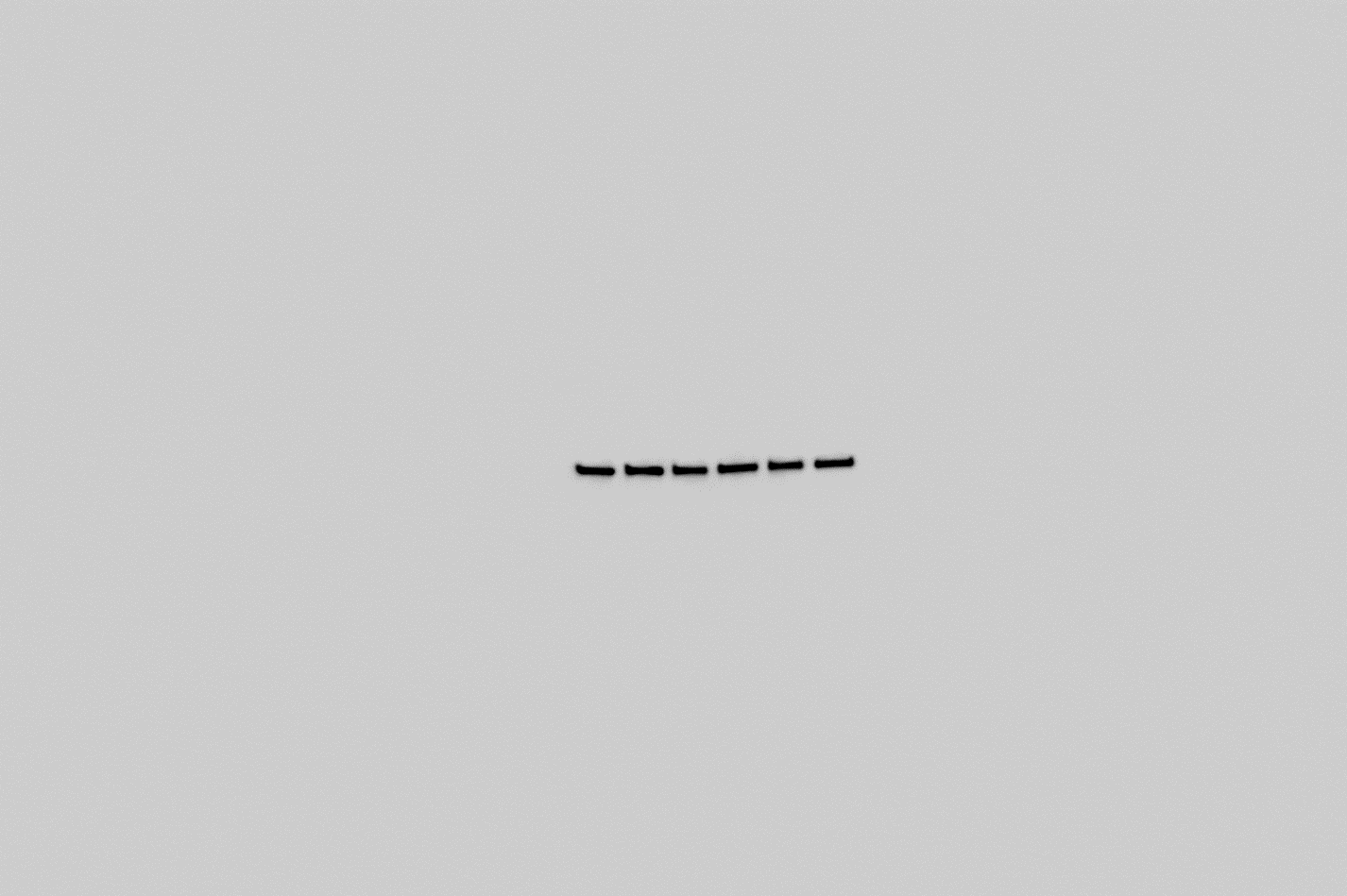

### Figure S2.

Raw images for CD68 Western blot samples (left three: MG single; right three: MG co)

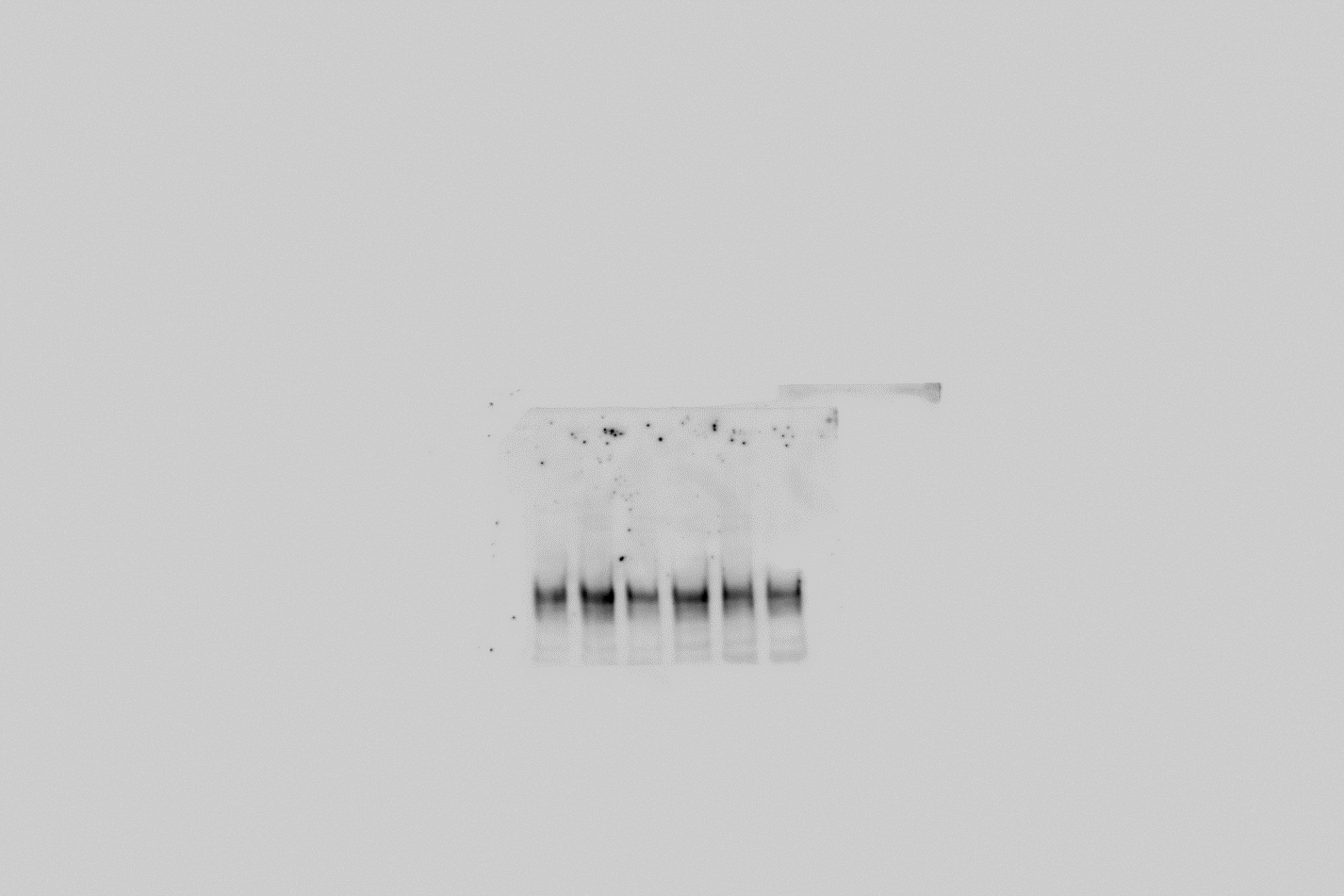

### Supplementary Tables

#### ***Table S1***.

Selected gene ontology (GO) biological process terms that showed significant differences in overrepresented test of GBM co-culture compared to single culture.

| **GO Term ID** | **P.Down** | **Term** |
| --- | --- | --- |
| GO:0050896 | 2.05E-16 | response to stimulus |
| GO:0030154 | 4.02E-13 | cell differentiation |
| GO:0051716 | 4.22E-13 | cellular response to stimulus |
| GO:0042127 | 3.18E-11 | regulation of cell proliferation |
| GO:0022610 | 1.43E-10 | biological adhesion |
| GO:0042221 | 2.55E-10 | response to chemical |
| GO:0007268 | 3.10E-10 | chemical synaptic transmission |
| GO:0007155 | 3.23E-10 | cell adhesion |
| GO:0016477 | 7.51E-10 | cell migration |
| GO:0048870 | 2.83E-09 | cell motility |
| GO:0048584 | 4.21E-08 | positive regulation of response to stimulus |
| GO:0070887 | 4.46E-08 | cellular response to chemical stimulus |
| GO:0098609 | 4.93E-08 | cell-cell adhesion |
| GO:0035295 | 1.25E-07 | tube development |
| GO:0043491 | 8.48E-06 | protein kinase B signaling |
| GO:0070372 | 2.93E-05 | regulation of ERK1 and ERK2 cascade |
| GO:0070371 | 7.71E-05 | ERK1 and ERK2 cascade |
| GO:0043408 | 0.00015 | regulation of MAPK cascade |
| **GO Term ID** | **P.Up** | **Term** |
| GO:0016071 | 2.18E-22 | mRNA metabolic process |
| GO:0006396 | 2.24E-21 | RNA processing |
| GO:0008380 | 5.48E-21 | RNA splicing |
| GO:0000375 | 1.10E-19 | RNA splicing, via transesterification reactions |
| GO:0090304 | 1.20E-19 | nucleic acid metabolic process |
| GO:0007049 | 1.10E-18 | cell cycle |
| GO:0051276 | 4.46E-17 | chromosome organization |
| GO:0022402 | 5.03E-17 | cell cycle process |
| GO:0000278 | 1.07E-16 | mitotic cell cycle |
| GO:0006397 | 1.51E-16 | mRNA processing |
| GO:0007059 | 1.13E-15 | chromosome segregation |
| GO:0006139 | 1.89E-15 | nucleobase-containing compound metabolic process |
| GO:0034641 | 3.21E-15 | cellular nitrogen compound metabolic process |
| GO:0051726 | 3.44E-15 | regulation of cell cycle |
| GO:0006725 | 7.78E-15 | cellular aromatic compound metabolic process |
| GO:0046483 | 1.11E-14 | heterocycle metabolic process |
| GO:0010564 | 4.84E-14 | regulation of cell cycle process |
| GO:1903047 | 5.16E-14 | mitotic cell cycle process |
| GO:0016070 | 1.31E-13 | RNA metabolic process |
| GO:1901360 | 3.65E-13 | organic cyclic compound metabolic process |
| GO:0098813 | 5.30E-13 | nuclear chromosome segregation |
| GO:0006259 | 8.83E-13 | DNA metabolic process |
| GO:1903311 | 9.38E-13 | regulation of mRNA metabolic process |
| GO:0044770 | 1.02E-12 | cell cycle phase transition |

#### Table S2.

Kyoto Encyclopedia of Genes and Genomes (KEGG) pathways that showed significant differences in overrepresented test of GBM co-culture compared to single culture.

| **KEGG ID** | **P.Down** | **avg.logfc.dir** | **NumGenes** | **GeneSet** |
| --- | --- | --- | --- | --- |
| hsa04640 | 1.15E-07 | -2.73974 | 17/97 | Hematopoietic cell lineage |
| hsa04060 | 2.54E-12 | -2.70804 | 43/294 | Cytokine-cytokine receptor interaction |
| hsa04610 | 4.59E-08 | -2.62891 | 18/79 | Complement and coagulation cascades |
| hsa04657 | 4.40E-09 | -2.61813 | 26/93 | IL-17 signaling pathway |
| hsa05134 | 1.29E-08 | -2.49458 | 21/55 | Legionellosis |
| hsa05332 | 8.94E-08 | -2.45877 | 10/41 | Graft-versus-host disease |
| hsa05133 | 7.74E-09 | -2.38943 | 24/76 | Pertussis |
| hsa04064 | 2.35E-11 | -2.36912 | 30/100 | NF-kappa B signaling pathway |
| hsa05164 | 1.30E-17 | -2.35976 | 50/167 | Influenza A |
| hsa04621 | 1.69E-17 | -2.26826 | 53/178 | NOD-like receptor signaling pathway |
| hsa04622 | 7.39E-08 | -2.2217 | 21/70 | RIG-I-like receptor signaling pathway |
| hsa04668 | 2.98E-16 | -2.18644 | 43/112 | TNF signaling pathway |
| hsa04620 | 2.32E-08 | -2.12804 | 27/104 | Toll-like receptor signaling pathway |
| hsa05160 | 3.55E-08 | -2.11714 | 36/155 | Hepatitis C |
| hsa05162 | 4.91E-10 | -2.11135 | 37/138 | Measles |
| hsa05169 | 6.20E-11 | -1.81214 | 48/201 | Epstein-Barr virus infection |
| hsa05167 | 1.33E-10 | -1.77906 | 46/186 | Kaposi's sarcoma-associated herpesvirus infection |
| hsa04380 | 4.02E-10 | -1.5421 | 33/128 | Osteoclast differentiation |
| hsa04010 | 4.38E-11 | -1.1984 | 63/295 | MAPK signaling pathway |
| hsa04510 | 8.18E-08 | -1.13458 | 45/199 | Focal adhesion |
| **KEGG ID** | **P.Up** | **avg.logfc.dir** | **NumGenes** | **GeneSet** |
| hsa04974 | 0.0218 | 1.86449 | 10/90 | Protein digestion and absorption |
| hsa04614 | 0.01741 | 1.29397 | 4/23 | Renin-angiotensin system |
| hsa04960 | 0.00344 | 1.12132 | 8/37 | Aldosterone-regulated sodium reabsorption |
| hsa04919 | 0.01225 | 1.09803 | 18/119 | Thyroid hormone signaling pathway |
| hsa04360 | 0.01229 | 1.07042 | 25/181 | Axon guidance |
| hsa04930 | 9.09E-04 | 1.04501 | 10/46 | Type II diabetes mellitus |
| hsa04973 | 0.002 | 1.03471 | 8/44 | Carbohydrate digestion and absorption |
| hsa04550 | 0.00275 | 0.92302 | 21/140 | Signaling pathways regulating pluripotency of stem cells |
| hsa00650 | 0.003 | 0.90608 | 6/28 | Butanoate metabolism |
| hsa04630 | 0.01063 | 0.78548 | 17/162 | JAK-STAT signaling pathway |
| hsa00600 | 6.10E-04 | 0.76163 | 11/47 | Sphingolipid metabolism |
| hsa05230 | 0.0084 | 0.75088 | 12/69 | Central carbon metabolism in cancer |
| hsa05213 | 0.01658 | 0.73896 | 11/58 | Endometrial cancer |
| hsa01521 | 0.00255 | 0.73389 | 16/79 | EGFR tyrosine kinase inhibitor resistance |
| hsa04213 | 0.0018 | 0.72778 | 13/62 | Longevity regulating pathway - multiple species |
| hsa04211 | 0.00336 | 0.71155 | 16/89 | Longevity regulating pathway |
| hsa04068 | 0.00117 | 0.70494 | 23/132 | FOXO signaling pathway |
| hsa00280 | 0.00193 | 0.68399 | 11/48 | Valine, leucine and isoleucine degradation |
| hsa04152 | 3.27E-03 | 0.68326 | 19/120 | AMPK signaling pathway |
| hsa04140 | 0.02299 | 0.52513 | 19/128 | Autophagy - animal |

#### Table S3.

Secretome profiling pixel intensities for individual blots.

|  | **1,2** | **3,4** | **5,6** | **7,8** | **9,10** | **11,12** | **13,14** | **15,16** | **17,18** | **19,20** | **21,22** | **23,24** |
| --- | --- | --- | --- | --- | --- | --- | --- | --- | --- | --- | --- | --- |
| **A** | Reference |  | Activin A | ADAMTS-1 | ANG | Ang-1 | Ang-2 | Angiostatin | Amphiregulin | Artemin |  | Reference |
| **B** | Coagulation Factor III | CXCL16 | DPPIV/CD26 | EGF | EG-VEGF / PK1 | Endoglin/ CD105 | Endostatin/ Collagen XVIII | Endothelin-1/ ET-1 | FGF acidic / FGF-1 | FGF basic/ FGFR2 | FGF-4 | FGF-7/ KGF |
| **C** | GDNF | GM-CSF | HB-EGF | HGF | IGFBP-1 | IGFBP-2 | IGFBP-3 | IL-1β/ IL-1F2 | IL-8/ CXCL8 | LAP (TGF-β1) | Leptin | MCP-1/ CCL2 |
| **D** | MIP-1α/ CCL3 | MMP-8 | MMP-9 | NRG1-β1/ HRG1-β1 | Pentraxin 3 (PTX3)/ TSG-14 | PD-ECGF | PDGF-AA | PDGF-AB/PDGF-BB | Persephin | Platelet Factor 4 (PF4)/ CXCL4 | PIGF | Prolactin |
| **E** | Serpin B5/Maspin | Serpin E1/ PIA-1 | Serpin F1/ PEDF | TIMP-1 | TIMP-4 | Thrombospondin-1/ TSP-1 | Thrombospondin-2/ TSP-2 | uPA | Vasohibin | VEGF | VEGF-C |  |
| **F** | Reference |  |  |  |  |  |  |  |  |  |  | Reference(-) |

| **Co-culture** | **1,2** | **3,4** | **5,6** | **7,8** | **9,10** | **11,12** | **13,14** | **15,16** | **17,18** | **19,20** | **21,22** | **23,24** |
| --- | --- | --- | --- | --- | --- | --- | --- | --- | --- | --- | --- | --- |
| **A** | 164084 |  | 5933 | 4511 | 77221 | 12324 | 30943 | 4382 | 10237 | 13623 |  | 172265 |
| **B** | 21127 | 100397 | 105680 | 2793 | 7415 | 9109 | 56740 | 58945 | 9621 | 11653 | 3666 | 10655 |
| **C** | 22080 | 159180 | 27719 | 6279 | 20742 | 117141 | 47737 | 21243 | 103269 | 5619 | 7779 | 193483 |
| **D** | 76160 | 23211 | 23510 | 9121 | 125618 | 8559 | 39272 | 5398 | 14324 | 6179 | 73447 | 14133 |
| **E** | 8021 | 142679 | 37367 | 151371 | 26849 | 118570 | 9063 | 149655 | 9145 | 146395 | 7705 |  |
| **F** | 164440 |  |  |  |  |  |  |  |  |  |  |  |
| **GBM12 single** | **1,2** | **3,4** | **5,6** | **7,8** | **9,10** | **11,12** | **13,14** | **15,16** | **17,18** | **19,20** | **21,22** | **23,24** |
| **A** | 171711 |  | 5241 | 4814 | 16742 | 10525 | 36898 | 3707 | 4616 | 13563 |  | 174334 |
| **B** | 15869 | 20244 | 37101 | 2084 | 5854 | 6876 | 13715 | 42223 | 5613 | 5792 | 2383 | 3098 |
| **C** | 5503 | 5140 | 8702 | 4726 | 13117 | 116702 | 21424 | 4637 | 6466 | 2803 | 3780 | 62826 |
| **D** | 12014 | 8573 | 12945 | 6719 | 51279 | 6579 | 4170 | 2530 | 10872 | 4698 | 8460 | 6173 |
| **E** | 6921 | 161090 | 20981 | 168026 | 28475 | 21582 | 8442 | 92865 | 7087 | 106160 | 4682 |  |
| **F** | 161378 |  |  |  |  |  |  |  |  |  |  |  |
| **MG single** | **1,2** | **3,4** | **5,6** | **7,8** | **9,10** | **11,12** | **13,14** | **15,16** | **17,18** | **19,20** | **21,22** | **23,24** |
| **A** | 168712 |  | 5285 | 3558 | 47268 | 10322 | 34060 | 3859 | 11917 | 14072 |  | 152739 |
| **B** | 12840 | 111061 | 83736 | 1825 | 6611 | 4315 | 56972 | 57779 | 7042 | 7427 | 1898 | 3370 |
| **C** | 19843 | 171287 | 19797 | 4258 | 13037 | 31342 | 15741 | 17478 | 118374 | 3590 | 3791 | 112104 |
| **D** | 41847 | 17375 | 19673 | 6807 | 128386 | 8996 | 68664 | 10749 | 12438 | 5068 | 55268 | 7013 |
| **E** | 7428 | 154884 | 23974 | 163052 | 8570 | 126200 | 10069 | 177304 | 9770 | 131554 | 5543 |  |
| **F** | 165137 |  |  |  |  |  |  |  |  |  |  |  |
| **Mix** | **1,2** | **3,4** | **5,6** | **7,8** | **9,10** | **11,12** | **13,14** | **15,16** | **17,18** | **19,20** | **21,22** | **23,24** |
| **A** | 183765 |  | 5681 | 4570 | 32750 | 10303 | 36857 | 4000 | 9825 | 14118 |  | 184350 |
| **B** | 14704 | 87033 | 57097 | 1812 | 6586 | 6192 | 43281 | 50330 | 8980 | 7451 | 2418 | 4614 |
| **C** | 10725 | 150283 | 14323 | 4587 | 17893 | 118573 | 18747 | 11414 | 141744 | 4774 | 4550 | 100233 |
| **D** | 13829 | 16149 | 17374 | 6308 | 124341 | 11577 | 40357 | 5998 | 16923 | 6672 | 34010 | 7706 |
| **E** | 9754 | 169931 | 23499 | 155625 | 24689 | 130373 | 11382 | 167493 | 10051 | 129604 | 5125 |  |
| **F** | 161503 |  |  |  |  |  |  |  |  |  |  |  |
